# Revealing the threat of emerging SARS-CoV-2 mutations to antibody therapies

**DOI:** 10.1101/2021.04.12.439473

**Authors:** Jiahui Chen, Kaifu Gao, Rui Wang, Guo-Wei Wei

## Abstract

The ongoing massive vaccination and the development of effective intervention offer the long-awaited hope to end the global rage of the COVID-19 pandemic. However, the rapidly growing SARS-CoV-2 variants might compromise existing vaccines and monoclonal antibody (mAb) therapies. Although there are valuable experimental studies about the potential threats from emerging variants, the results are limited to a handful of mutations and Eli Lilly and Regeneron mAbs. The potential threats from frequently occurring mutations on the SARS-CoV-2 spike (S) protein receptor-binding domain (RBD) to many mAbs in clinical trials are largely unknown. We fill the gap by developing a topology-based deep learning strategy that is validated with tens of thousands of experimental data points. We analyze 261,348 genome isolates from patients to identify 514 non-degenerate RBD mutations and investigate their impacts on 16 mAbs in clinical trials. Our findings, which are highly consistent with existing experimental results about variants from the UK, South Africa, Brazil, US-California, and Mexico shed light on potential threats of 95 high-frequency mutations to mAbs not only from Eli Lilly and Regeneron but also from Celltrion and Rockefeller University that are in clinical trials. We unveil, for the first time, that high-frequency mutations R346K/S, N439K, G446V, L455F, V483F/A, E484Q/V/A/G/D, F486L, F490L/V/S, Q493L, and S494P/L might compromise some of mAbs in clinical trials. Our study gives rise to a general perspective about how mutations will affect current vaccines.

## 1 Introduction

Since the first positive cases of coronavirus disease, 2019 (COVID-19) caused by severe acute respiratory syndrome coronavirus 2 (SARS-CoV-2) was reported in late December 2019, over 2.5 million lives have been taken away in the COVID-19 pandemic up to March 10, 2021. The developments of vaccines and antibody therapies are the most significant scientific accomplishments that offer the essential hope to win the battle against COVID-19. Nonetheless, the emerging SARS-CoV-2 variants signal a major threat to existing vaccines and antibody drugs.

SARS-CoV-2 is a novel *β*-coronavirus, which is an enveloped, unsegmented positive-sense single-strand ribonucleic acid (RNA) virus. It gains entry into the host cell through the binding of its spike (S) protein receptor-binding domain (RBD) to the host angiotensin-converting enzyme 2 (ACE2) receptor, primed by host transmembrane protease, serine 2 (TMPRSS2) [1]. According to epidemiological and biochemical studies, the binding free energy (BFE) between the S protein and ACE2 is proportional to the infectivity of different SARS-CoV-2 variants in the host cells [2,3]. Intrinsically, the mutation-induced BFE changes (ΔΔ*G*) of S protein and ACE2 complex provide a method to measure the infectivity changes of a SARS-CoV-2 variant compared to the first SARS-CoV-2 strain that deposited to GenBank (Access number: NC 045512.2) [4]. Specifically, the positive mutation-induced BFE change of S and ACE2 indicates that this mutation would strengthen the infectivity of SARS-CoV-2, while the negative mutation-induced BFE change reveals the possibility of the weakening transmissible and infectious. Therefore, one can predict the impact of SARS-CoV-2 RBD variants on infectivity by estimating their BFE changes [4–6].

Moreover, the binding of S protein and ACE2 will trigger the host adaptive immune system to produce antibodies against the invading virus [7,8]. As illustrated in Figure 1, antibodies are secreted by a type of white blood cell called B cell (mainly by plasma B cells or memory B cells). An antibody can either attach to the surface of B cell (called B-cell receptor (BCR)) or exist in the blood plasma in a solute form. An antibody can be generated in three ways: 1) Once SARS-CoV-2 invades the host cell, the adaptive immune system will be triggered, and the B cells will generate and secrete antibodies. 2) In antibody therapies, antibodies are initially generated from patient immune response and T-cell pathway inhibitors [7], which are called antibody drugs [8]. Most COVID-19 antibody drugs primarily target S protein. 3) The vaccine is designed to stimulate an effective host immune response, which is another way to make B cells secrete antibodies [9]. At this stage, various vaccines, including two mRNA vaccines designed by Pfizer-BioNTech and Moderna, have been granted authorization for emergency use in many countries, aiming to give our human cells instructions to make a harmless S protein piece to initiate the immune response actively. Although COVID-19 vaccines are the gamechanger, S protein mutations might weaken the binding between the SARS-CoV-2 S protein and antibodies and thus, reduce the efficiency and efficacy of the existing vaccines and antibody therapies [10].

**Figure 1:**
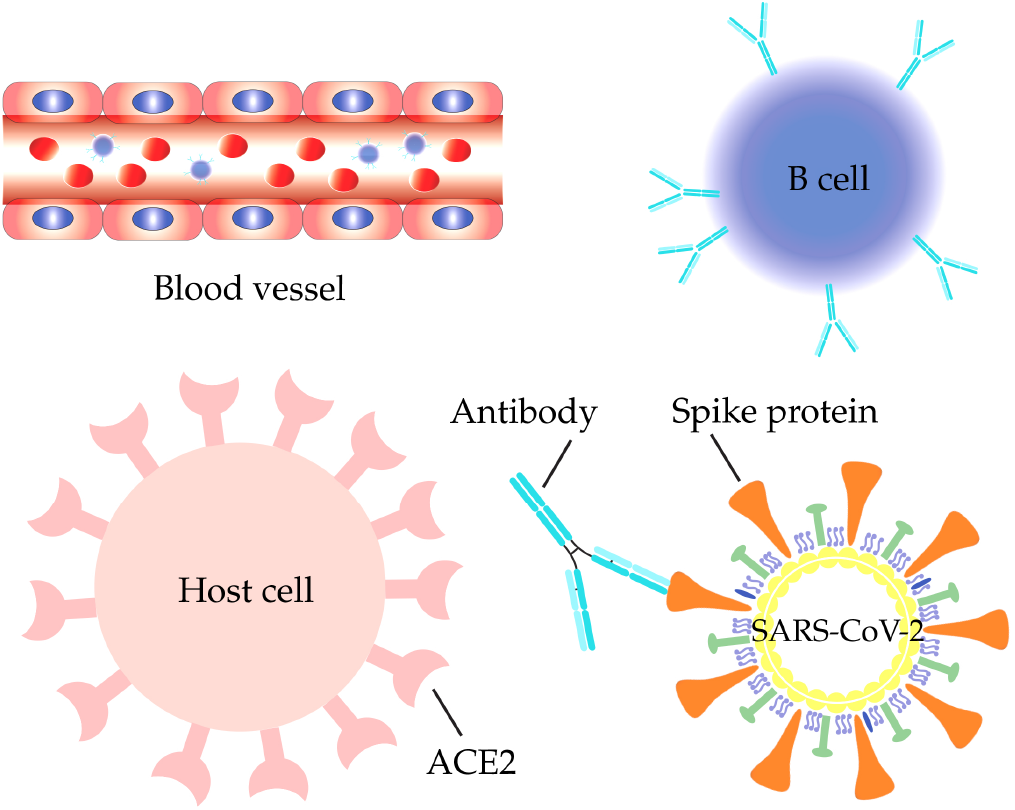
SARS-CoV-2 S protein antibodies are secreted by B cells in aiming to compete with the host ACE2 for binding to the S protein RBD.

Although SARS-CoV-2 has a higher fidelity in the replication process which benefits from its genetic proofreading mechanism regulated by the non-structural protein 14 (NSP14) and RNA-dependent RNA polymerase (RdRp) [11, 12], over 5,000 unique mutations has been found on SARS-CoV-2 S protein [5], which raises the question that how these mutations on S protein will affect the existing vaccines and antibody drugs. Antibody resistance of SARS-CoV-2 variants B.1.351 and B.1.1.7 was reported [10]. Mutation E484K on S protein RBD may help SARS-CoV-2 slip past the host immune defenses, is broadly founded in the B.1.351 (a.k.a 20H/501Y.V2) variant [13] and the P.1 (a.k.a 20J/501Y.V3) variant [14]. The ongoing evaluation of susceptibility of variants in subjects treated with the antibody-drug bamlanivimab shows that E484K substitution in B.1.1.7, P.1, and B.1.526 variants had reduced susceptibility to bamlanivimab [15]. Moreover, the K417N+E484K+N501Y substitutions in B.1.351 and P.1 variants had also reduced susceptibility to bamlanivimab [15]. Specifically, a 50% increment in the transmission of the B.1.351 variant is estimated in [16]. Both P.1 and B.1.351 variants cause negative effects on the neutralization by emergency use authorization (EUA) monoclonal antibody therapeutics [17, 18], and the moderate reductions in neutralizing activity were observed by using convalescent and post-vaccination sera [19]. Furthermore, the B.1.427/B.1.429 variant that is initially found in California carries an L452R mutation on the S protein RBD, which approximately increase 20% of the transmissibility of SARS-CoV-2 [19], and has a mild negative impact on neutralization by some EUA therapeutics according to Food and Drug Administration (FDA) report [15, 20]. Notably, by using convalescent and post-vaccination sera, moderate reductions in neutralizing activity of L452R were observed [19].

However, the determination of whether a mutation will reduce susceptibility to the existing antibodies and antibody drugs from the wet laboratory experiments is time-consuming. Current experimental studies are restricted to only a small fraction of known RBD mutations that have been observed. There is no reliable measurement about whether a mutation will evade a vaccine because none knows how many different antibodies will be created from the vaccination. Based on the molecular mechanism of SARS-CoV-2 infectivity, antibody, and vaccine, one can quantitatively estimate mutation impacts on SARS-CoV-2 infectivity and an antibody-drug through computing mutation-induced BFE changes of the S protein-ACE2 complex and the S protein-antibody complex, respectively. In our earlier work, we proposed a TopNetTree model to predict the RBD-induced binding free energy (BFE) changes of S protein with ACE2 and 56 antibodies [5]. We showed that RBD mutation N501Y can significantly strengthen SARS-CoV-2 infectivity [5], which is consistent with experiment [16]. Our results indicated that K417N, E484K, and L452R are all the antibody-escape mutations with positive BFE changes, which are consistent with the findings from many wet labs [5,10]. Among them, mutation L452R in the California variant is both more infectious and antibody resistant [5]. We found that the T478K mutation in Mexico variant (B.1.1.222) has the most significant high value of predicted BFE changes and is one of the potential vaccine-escape [5], which has a rapid growth rate in Mexico. Our prediction is confirmed from a report that mutation T478K is spreading at an alarming speed [21]. We also predicted 1149 most likely, 1912 likely, and 625 unlikely receptor-binding domain (RBD) mutations [6]. Currently, all known RBD mutations were correctly predicted as the most likely ones in our work [5, 6].

The objective of this work is to reveal the mutational threats to 16 antibody drug candidates that are either in clinical trials or associated with clinical trial antibodies as shown in Figure 2. To this end, we analyze 261,348 complete SARS-CoV-2 genome sequences isolated from patients to identify 27,530 unique single mutations. Among them, 514 non-degenerate mutations are found on the S protein RBD of SARS-CoV-2. We develop a deep learning model-based algebraic topology to estimate the mutation-induced BFE changes. We show that 95 RBD mutations that have been observed over 10 times have favorable predicted BFE changes, which further confirms the accuracy and reliability of our predictions from the epidemiological point of view. Our study of antibody drug candidates is valuable and complementary to experimental results in the following senses. First, our machine learning and deep learning models validated with tens of thousands of experimental data points, including SARS-CoV-2 related deep mutations, are reliable as confirmed by emerging experimental data on various SARS-CoV-2 variants. Second, there are 95 fast-growing RBD mutations around the world that pose imminent threats to existing and future vaccines and antibody therapies. The current experimental capability lags behind the rapidly growing RBD mutations. For example, there is no experimental study about the rapidly increasing Mexico variant B.1.1.222. Our approach helps close the gap. This work provides a threat analysis of all 514 existing RBD mutations. However, our emphasis is given to 95 fast-growing RBD mutations. Third, the current experiments in the literature are limited to two EUA monoclonal antibody therapeutics from Regeneron [22] and Eli Lilly [8]. We extend our analysis to many other antibody therapeutic candidates that are in various stages of clinical trials, such as those from Celltrion [23] and the Rockefeller University.

**Figure 2:**
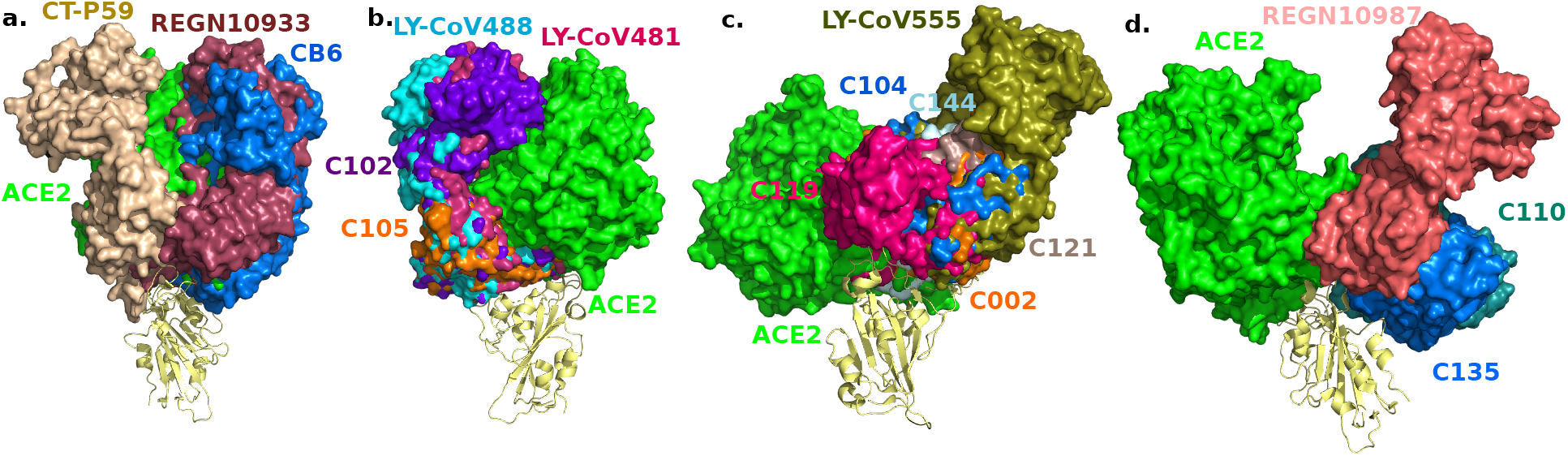
3D alignment of 16 antibodies and ACE2 on the S protein RBD. a. CT-P59 (7CM4), REGN10933 (6XDG), CB6 (7C01). b. LY-CoV488 (7KMH), LY-CoV481 (7KMI), C102 (7K8M), C105 (6XCM). c. LY-CoV555 (7KMG), C002 (7K8T), C104 (7K8U), C119 (7K8W), C121 (7K8X), C144 (7K90). d. REGN10987 (6XDG), C110 (7K8V), C135 (7K8Z).

## 2 Results

### 2.1 Antibodies in clinical trials

In this work, we study 16 antibodies, including 5 antibodies in phase 3 clinical trials or EUA, and 2 antibodies in phase 1 clinical trials. The rest of antibodies are closely related to those in clinical trials. For the 5 antibodies in phase 3 clinical trials or EUA, there are two antibody combination treatments, casirivimab/imdevimab (REGN10933/REGN10987), and bamlanivimab/etesevimab (LY-CoV555/CB6), and one single antibody treatment, regdanvimab (CT-P59) from Celltrion. C135 and C144 are two antibodies from the Rockefeller University in phase 1 clinical trials. The rest antibodies are C102, C105, C002, C104, C110, C119, C121, LY-CoV481, and LY-CoV488. Most of the antibodies are isolated or derived from COVID-19 human neutralizing antibodies [23–27], while REGN10933 and REGN10987 are derived from the treatments for Ebola – one from humanized mice and one from a convalescent patient [22]. According to the literature [22, 24, 25], antibodies REGN10933, REGN10987, LY-CoV555, and CB6, were optimized through fluorescence-activated cell sorters.

In Figure 2, we align 16 three-dimensional (3D) antibody structures with ACE2. Figures 2 a and b show 7 antibodies that directly compete with ACE2 on the binding domain. Three clinical-trial antibodies, namely CT-P59, REGN10933, and CB6, can be found in Figure 2 a. Figure 2 c shows 6 antibodies whose binding domains partially overlap with that of ACE2. Among them, LY-CoV555 and C144 are in clinical trials. Figure 2 d shows 3 antibodies that partially share their binding domains with ACE2. Antibodies REGN10987 and C135 do not compete with ACE2 directly and thus, they can be complements of other antibodies.

### 2.2 Impacts of SARS-CoV-2 on antibody efficacy and infectivity

SARS-CoV-2 variants with specific genetic markers are correlated to BFE changes on the RBD, degrade the neutralization by antibody treatments, or antibodies of the self immune system, and increase the difficulty of virus diagnostic or transmissible prediction. Especially, for those mutations that enhance transmissibility and weaken antibody neutralization, they should be prioritized in the investigation. In Figure 3, we illustrate RBD mutations involved in the UK variant B.1.1.7, Brazil variant P.1, South Africa variant B.1.351, US-California variant B.1.427, and Mexico variant B.1.1.222. In this figure, each RBD residue is colored by the maximum mutation-induced BFE change on the S protein-ACE2 complex from 19 possible mutations. One can notice that all the six mutations in various variants have positive BFE changes that enhance the binding of S protein RBD and ACE2, and consequently, the infectivity of SARS-CoV-2.

**Figure 3:**
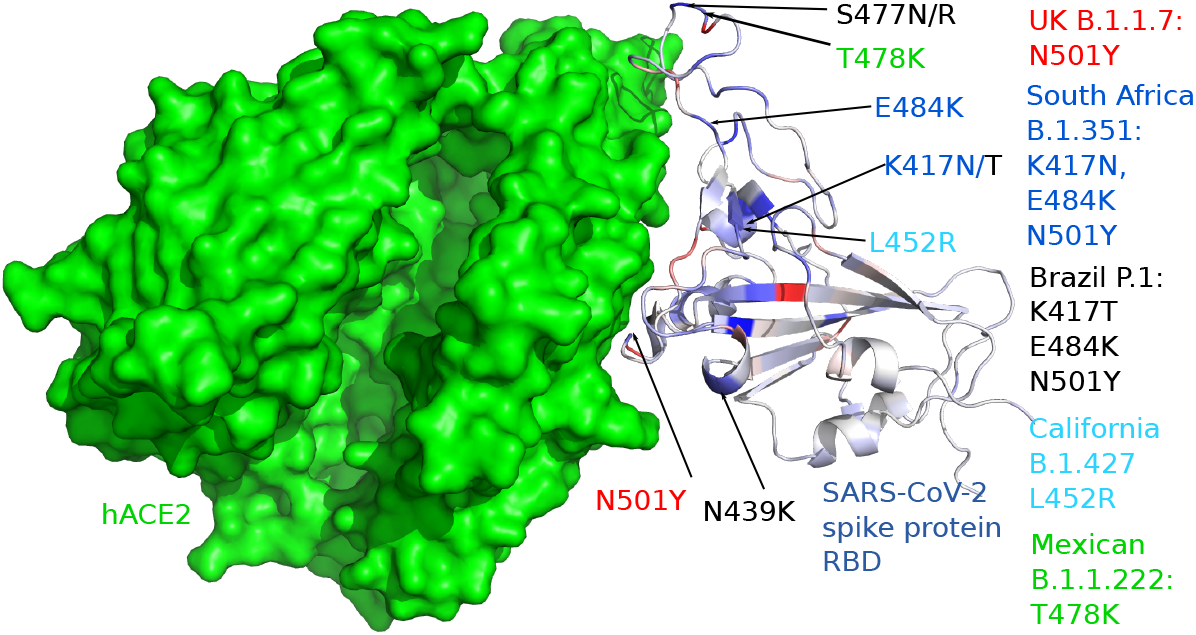
3D structure of human ACE2 (hACE2) and RBD. Color on the RBD structure indicates the BFE changes induced by mutations, where blue means binding strengthening and red means weakening.

Figure 4 gives an illustration of SARS-CoV-2 S protein RBD mutation-induced BFE changes to the complexes of S protein with antibodies or ACE2. Here, we only consider those mutations that have been observed more than 10 times, and a similar study for all known RBD mutations is presented in the Supporting information. Note that there is a strong correlation between the positive predicted mutation-induced BFE changes and observed mutation frequency. For a given mutation, if its BFE changes for antibodies are very negative value while for ACE2 very positive, then this mutation has a combined antibody-escape and fast-growing effort. Therefore, one can observe that mutations, R346K/S, K417T/N, L452R, E484K/Q, F486L, F490L/S, S484P/L, and N501Y, have this effect, while R346K/S and N501Y induce a relatively moderate weakening effect to most antibodies.

**Figure 4:**
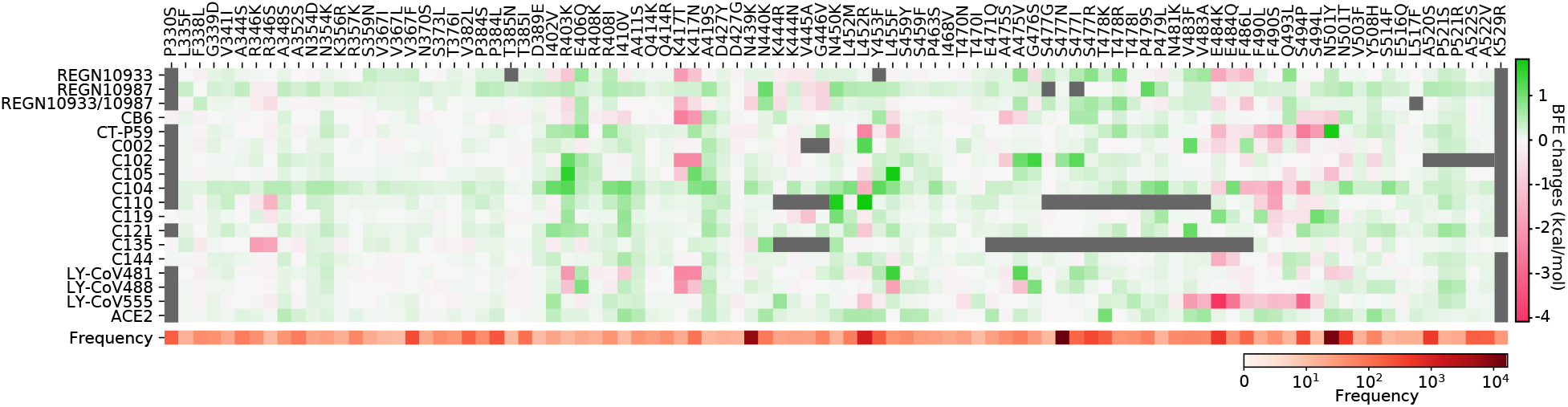
Illustration of the BFE changes of the complexes of S protein and antibodies or ACE2, induced by RBD mutations with frequencies being greater than 10. Positive changes strengthen the binding while negative changes weaken the binding. Here, only mutations that occurred on the relevant random coil of the S protein RBD are considered. The Grey color indicates that PDB structures do not involve specific residues.

Figure 5 shows the BFE changes induced by six RBD mutations for the S protein complexes with antibodies and ACE2. First of all, it is noted that all RBD mutations give positive BFE changes for binding to ACE2, leading to more infectious variants. Additionally, the magnitude of BFE changes on each mutation is correlated to the distance to antibodies. Therefore, antibodies having more overlap with ACE2 are impacted more significantly by mutations. For example, according to their 3D alignment in Figure 2, CB6, CT-P59, REGN10933, C102, C105, LY-CoV481, and LY-CoV488 who are directly competing with ACE2 have large BFE changes in five mutations. Antibodies that partially overlap with ACE2 in terms of binding domain, i.e., C002, C104, C119, C121, C144, and LY-CoV555, have only a few significant BFE changes. Antibodies, C110, C135, and REGN10987, which bind to the other side of the RBD, have very mild changes in all the mutations.

**Figure 5:**
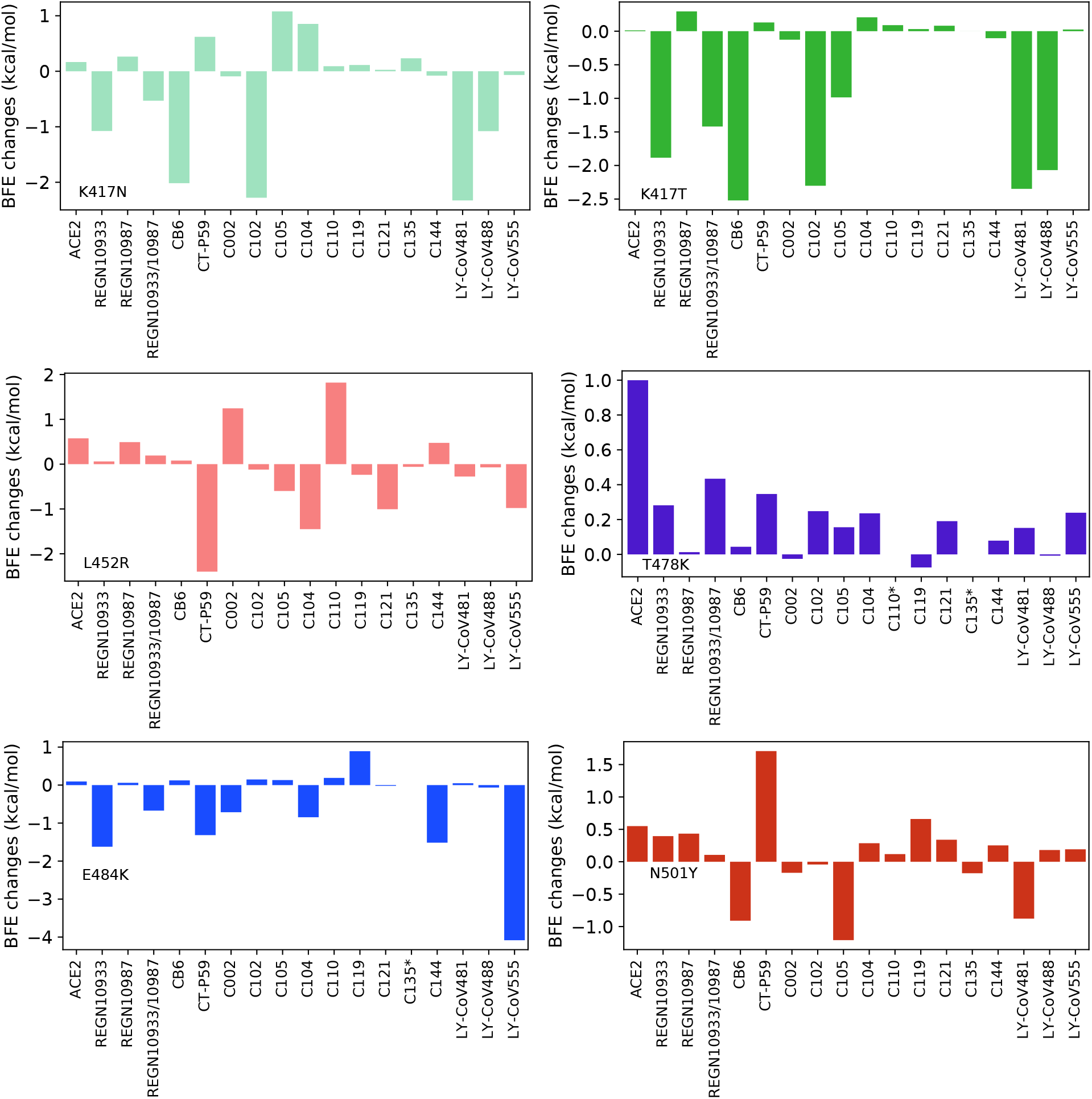
BFE changes induced by new SARS-CoV-2 mutations, K417N, K417T, L452R, T478K, E484K, and N501Y. C110* and C135*: no results due to incomplete PDB structure.

More specifically, mutation T478K whose frequency has risen exponentially since early 2021 in Mexico (Mexican B.1.1.222) induces a very large positive BFE change in the ACE2-S protein complex. This could explain why T478K is a fast-growing mutation although it might not affect the binding of antibodies to the S protein. As for three variants from UK (B.1.1.7), Brazil (P.1), and South Africa (B.1.351), they share the same mutation, N501Y, while the UK variant is the only one that contains one mutation on RBD and Brazil and South Africa variants contain other mutations K417N/T and E484K. Meanwhile, the experimental results show that most antibodies demonstrated neutralizing capability against the UK variant [17, 18, 28]. Interestingly, as reported by European Medicines Agency [29], regdanvimab (CT-P59) shows neutralizing ability against the UK variant. These results are highly consistent with the small positive BFE changes of N501Y on antibodies in Figure 5. For the key substitution, L452R, of California (B.1.427/429), regdanvimab (CT-P59), and bamlanivimab (LY-CoV555) have large negative BFE changes. In the FDA report of bamlanivimab (LY-CoV555) and etesevimab (CB6) [17, 18], the mutation L452R has a large fold reduction in susceptibility of single bamlanivimab and mild fold reduction of the combination of bamlanivimab and etesevimab. Lastly, we study the South Africa and Brazil variants, which share the same mutations E484K and N501Y but different in K417N/T. For antibodies in EUA, REGN10987 has mild changes on mutations, K417N/T and E484K, while REGN10933 and CB6 respond with large negative changes and LY-CoV555 has a significant negative change on E484K. Our predictions for the South Africa and Brazil variants are in excellent agreement with experimental data [28, 30].

### 2.3 Mutation impacts on antibodies in clinical trials

In this section, we study five antibodies in clinical trials or emergency use authorization. Two antibodies of Regeneron Pharmaceuticals, casirivimab and imdevimab, are studies together, followed by other three antibodies in phase 3, regdanvimab, bamlanivimab, and etesevimab. Two antibodies in phase 1 are discussed in the end as well. We emphasize those RBD mutations that have been observed for more than 10 times and denoted their as high-frequency mutations. A complete study of all known RBD mutations is given in the Supporting information.

#### 2.3.1 Antibodies REGN10933 and REGN10987 (aka Casirivimab and Imdevimab)

Regeneron’s Casirivimab and Imdevimab antibody cocktail against SARS-CoV-2 is the first combination therapy, which receives an FDA emergency use authorization. As the only one in the clinical trial antibodies that have the 3D structure of two antibodies binding to the RBD, we first study the BFE changes of them as an antibody combination. We examine the BFE changes induced by RDB mutations whose frequencies are greater than 10 in Figure 6 of the antibody cocktail, REGN10933 and REGN10987, binding to the S protein RBD. The single antibody analysis is provided in the Supporting information. Notice that mutations, K417T, N439K, G446V, E484K, E484G, and F486L, lead to large negative BFE changes. For positive BFE changes, it is good to see that there are high-frequency mutations, which indicates that this antibody combination potentially prevents the new variants of SARS-CoV-2, especially for variants with mutations L452R, S477N, and K501Y. However, some mutations with negative BFE changes have a very large magnitude, indicating that the antibody combination of REGN10933 and REGN10987 was an immune product optimized for the original un-mutated S protein. In general, parts of the mutations on the S protein weaken the REGN10933+REGN10987 binding and make the antibodies less competitive to ACE2. This cocktail is prone to South Africa and Brazil variants (K417N/T, E484K) but remains effective for UK and US-California variants (L452R and K501Y).

**Figure 6:**
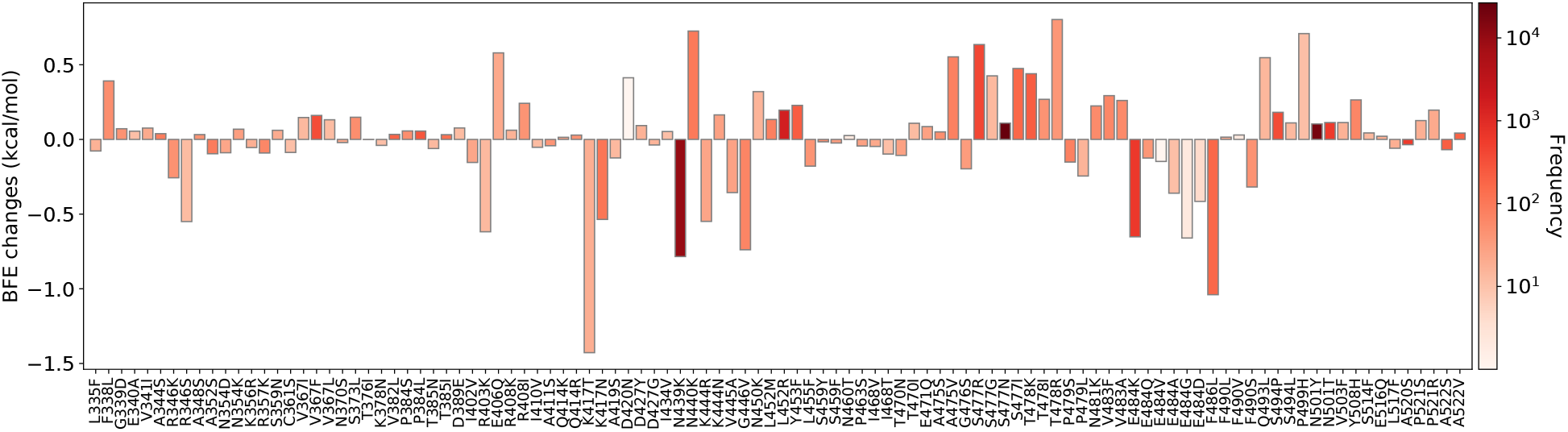
Illustration of SARS-CoV-2 mutation-induced binding free energy changes for the complexes of S protein and antibodies REGN10933 and REGN10987 (PDB: 6XDG). Here, mutations K417T, N439K, G446V, E484K/G, and F486L could potentially disrupt the binding of antibodies and S protein RBD.

**Figure 7:**
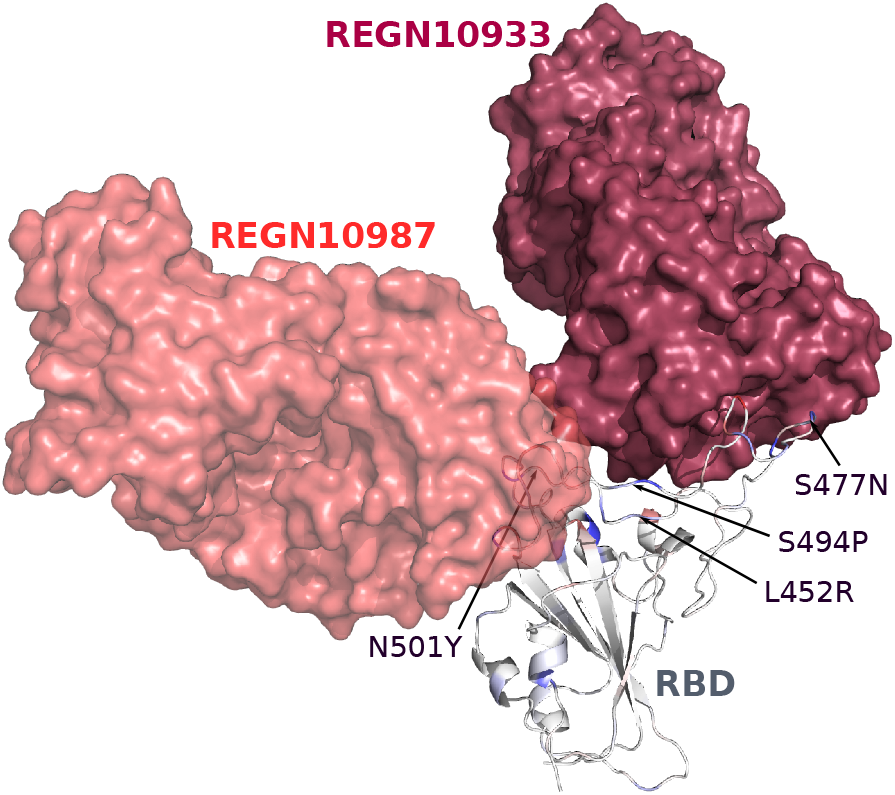
The binding complex of S protein RBD and REGN10933+REGN10987.

Additionally, these two antibodies can be studied separately as shown in Figures 8 and 9. By comparing the stand-alone BFE predictions to those in Figure 6, it can be concluded that antibody REGN10933 plays the main role in the antibody neutralization, while the antibody REGN10987 is a complement for two reasons. First, the antibody REGN10933 shares the same disrupted mutations with the combination and has larger BFE changes on those mutations. Secondly, the BFE changes for REGN10987 are mild, and most of them are positive values. According to the 3D alignment, the antibody REGN10987 does not directly compete with ACE2. Lastly, in the comparison, one can notice that the magnitude of BFE changes is smaller on the mutations for the combination. This indicates a more stable binding of the antibody combination.

**Figure 8:**
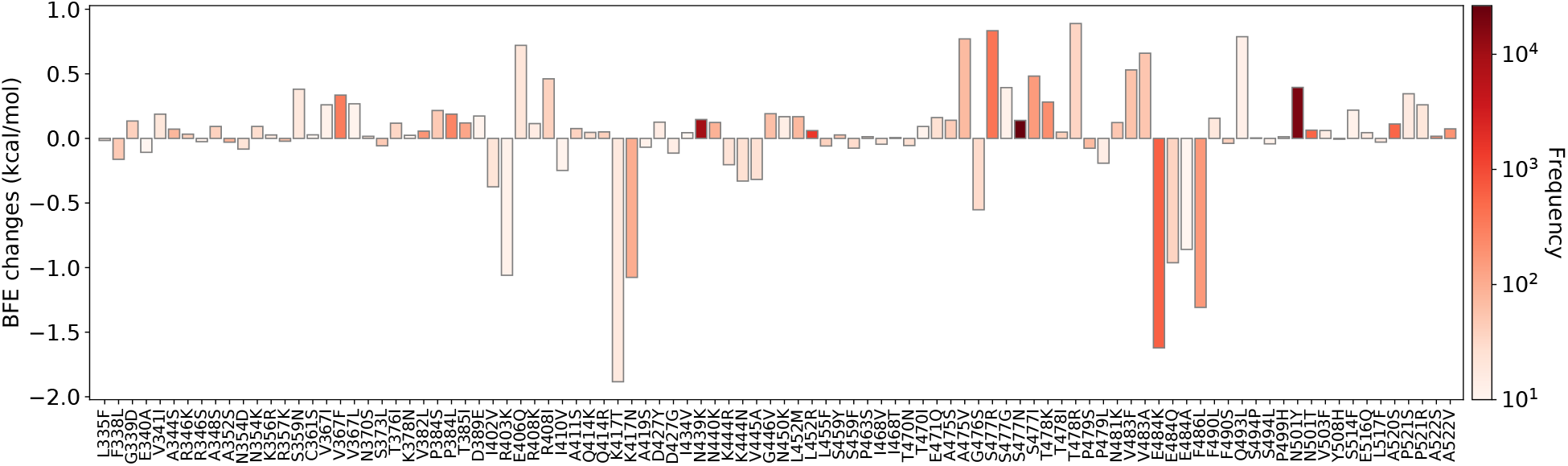
Illustration of SARS-CoV-2 mutation-induced binding free energy changes for the complexes of S protein and antibody REGN10933 (PDB: 6XDG). Here, mutations K417T, N439K, G446V, E484K/G, and F486L could potentially disrupt the binding of antibody and S protein RBD.

**Figure 9:**
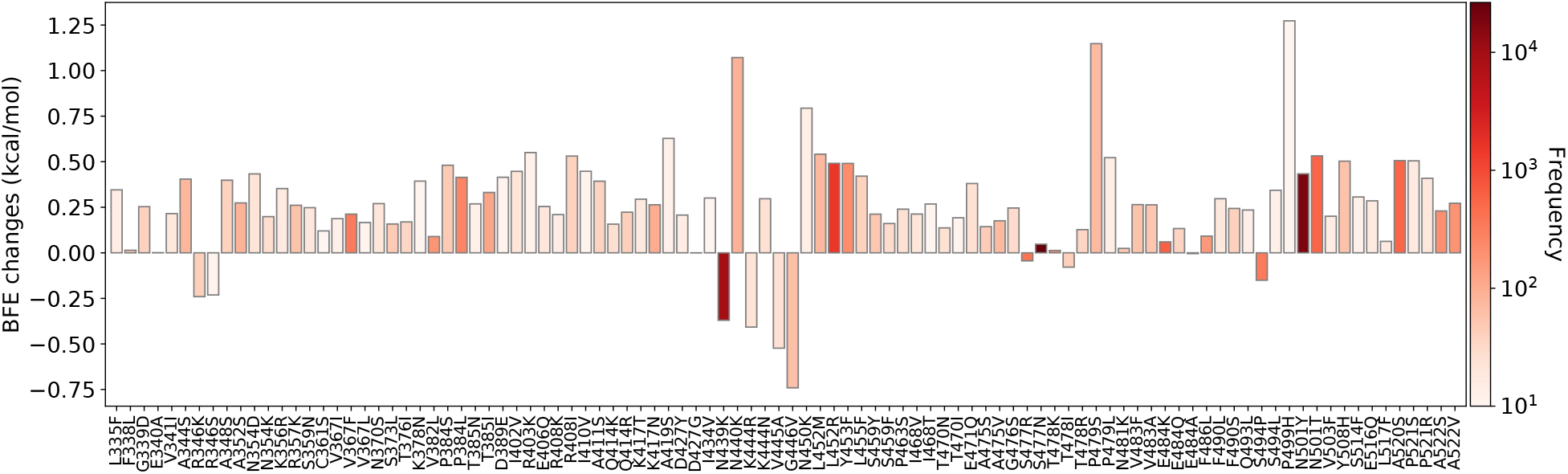
Illustration of SARS-CoV-2 mutation-induced binding free energy changes for the complexes of S protein and antibody REGN10987 (PDB: 6XDG).

#### 2.3.2 Antibodies LY-CoV555 and CB6 (aka Bamlanivimab and Etesevimab)

Bamlanivimab (LY-CoV555) was first developed as a single antibody therapy for the treatment of mild to moderate COVID-19 illness. However, it is not distributed alone due to the SARS-CoV-2 variant resistance and is used as an antibody combination with Etesevimab (CB6). Here, we first examine Bamlanivimab’s response to S protein RBD mutations followed by the discussion of Etesevimab.

**Figure 10:**
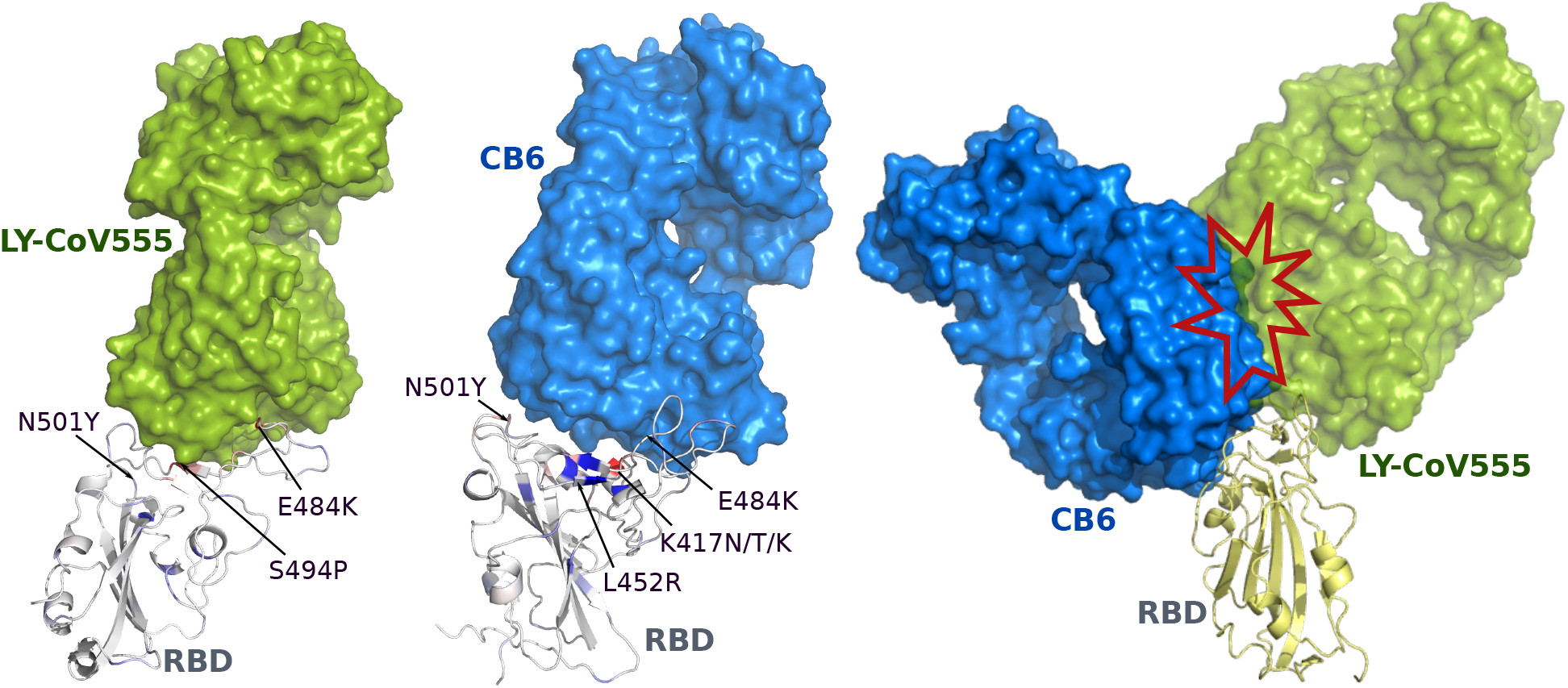
The binding complexes of S protein RBD with LY-CoV555 and CB6. There is a crash at the interface between two antibodies.

In the BFE changes prediction of LY-CoV555 (PDB: 7KMG) as shown in Figure 11, most mutations have mild changes, while mutations, L452R, V483F/A, E484K/Q/V/A/G/D, F486L, F490L/V/S, Q493L, and S494P, have large negative BFE changes. For positive BFE changes, the largest value is only 0.75 kcal/mol and the average of positive BFE changes is 0.16 kcal/mol. However, many mutations with negative BFE changes have very large magnitudes such that 7 mutations having binding free energy less than −2 kcal/mol, and the least value is −4.1 kcal/mol for E484K. This could indicate that antibody LY-CoV555 was an immune product optimized with respect to the original un-mutated S protein. In general, the mutations on S protein weaken the LY-CoV555 binding to S protein and make it less competitive with ACE2 as most mutations strengthen the S protein and ACE2 binding. The South Africa variant (E484K) and US-California variant (L452R) have a strong antibody-escape effect.

**Figure 11:**
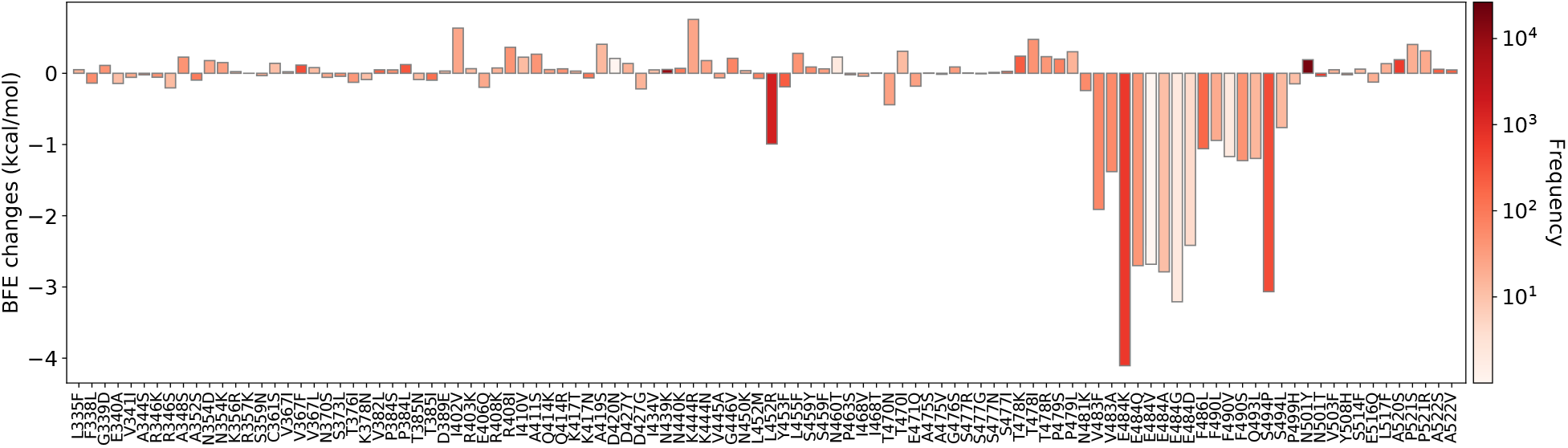
Illustration of SARS-CoV-2 RBD mutation-induced binding free energy changes for the complexes of S protein and antibody LY-CoV555 (PDB: 7KMG). Here, mutations L452R, V483F/A, E484K/Q/V/A/G/D, F486L, F490L/V/S, Q493L, and S494P could potentially disrupt the binding of antibodies and S protein RBD.

In Figure 12, we illustrate the mutation-induced BFE changes for antibody CB6 (PDB: 7C01), which directly competes with ACE2. One can notice that K417T, K417N, A475S, and N501Y, have large negative BFE changes, and three of them belong to SARS-CoV-2 variants. The rest mutations have a small magnitude of changes, and there are no large positive BFE changes. Antibody CB6 is isolated from peripheral blood mononuclear cells of patients convalescing from COVID-19 at the early stage and optimized based on an early version of the SARS-CoV-2 virus.

**Figure 12:**
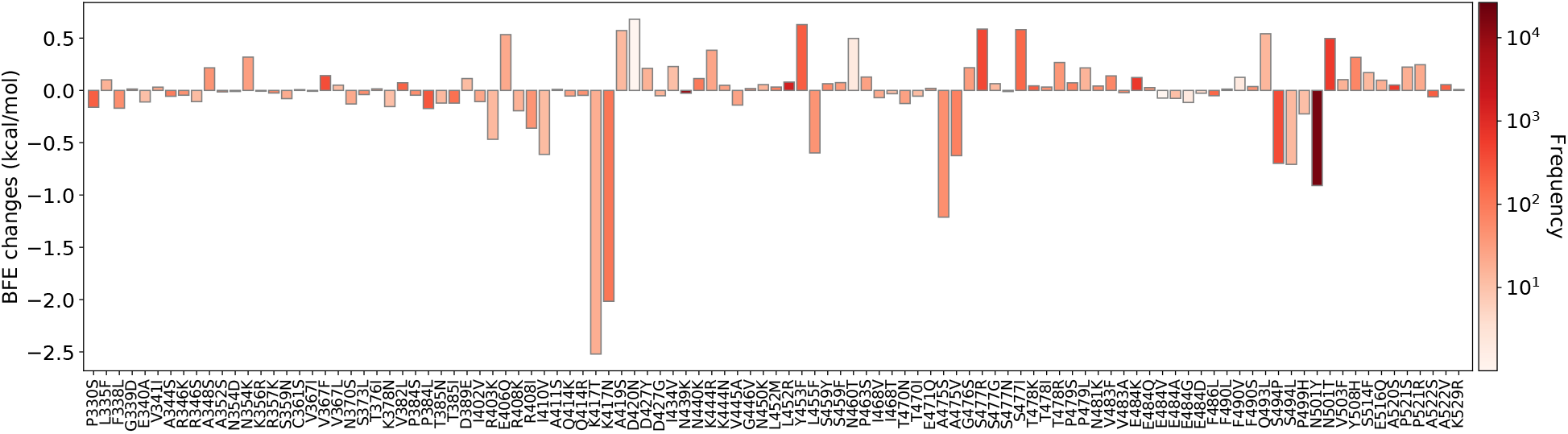
Illustration of SARS-CoV-2 mutation-induced binding free energy changes for the complexes of S protein and antibody CB6 (PDB: 7C01). Here, mutations K417T, N439K, G446V, E484K, E484G, and F486L could potentially disrupt the binding of antibody and S protein RBD.

In the 3D alignment, antibodies LY-CoV555 and CB6 share a partial binding domain with ACE2. Therefore, they are not only competing with ACE2 but also with each other. Comparing the BFE change prediction on both LY-CoV555 and CB6, one can note that two antibodies respond to S protein RBD mutations differently and thus are complementary. We deduce that the combined antibody will enhance the single antibody neutralization.

#### 2.3.3 Antibody CT-P59

Regdanvimab (CT-P59) has been approved for emergency use treatment in South Korea and is under review by European Medicines Agency (EMA). We present the BFE changes in Figure 14. Antibody CT-P59 shares a similar binding domain with ACE2 and is a potent candidate for the direct neutralization of SARS-CoV-2. Most mutations induce small changes in the binding free energy, while mutations L452R, L455F, E484K/A, F490L/S, and S494P/L induce large negative BFE changes. This indicates antibody CT-P59 has an antibody-escape effect for many variants, including the South Africa variant (B.1.351 with E484K) and the US-California (B.1.427 and B.1.429 with L452R). It is noticed that is that CT-P59 has a large positive BFE change for mutation N501Y, indicating CT-P59 can neutralize the SARS-CoV-2 UK variant (B.1.17).

**Figure 13:**
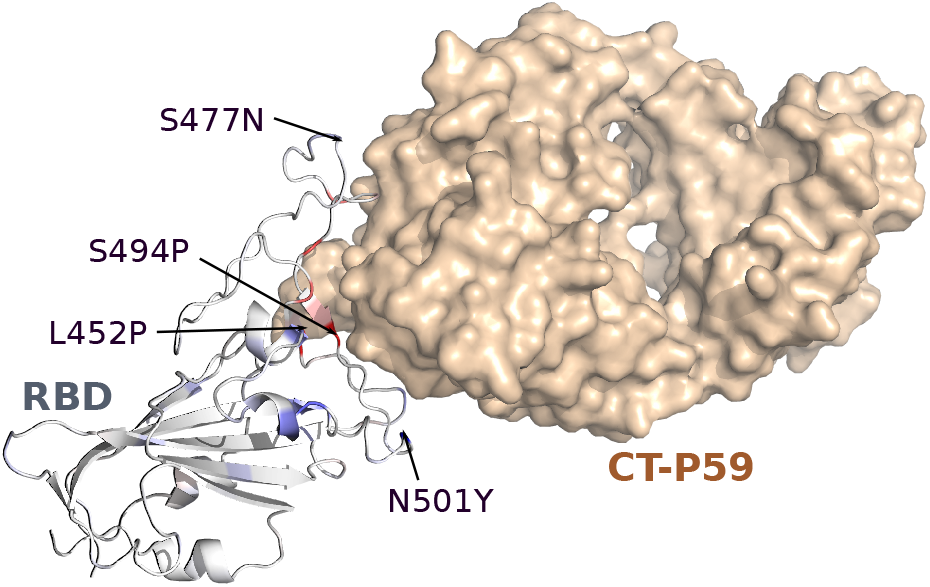
The binding complex of S protein and CT-P59.

**Figure 14:**
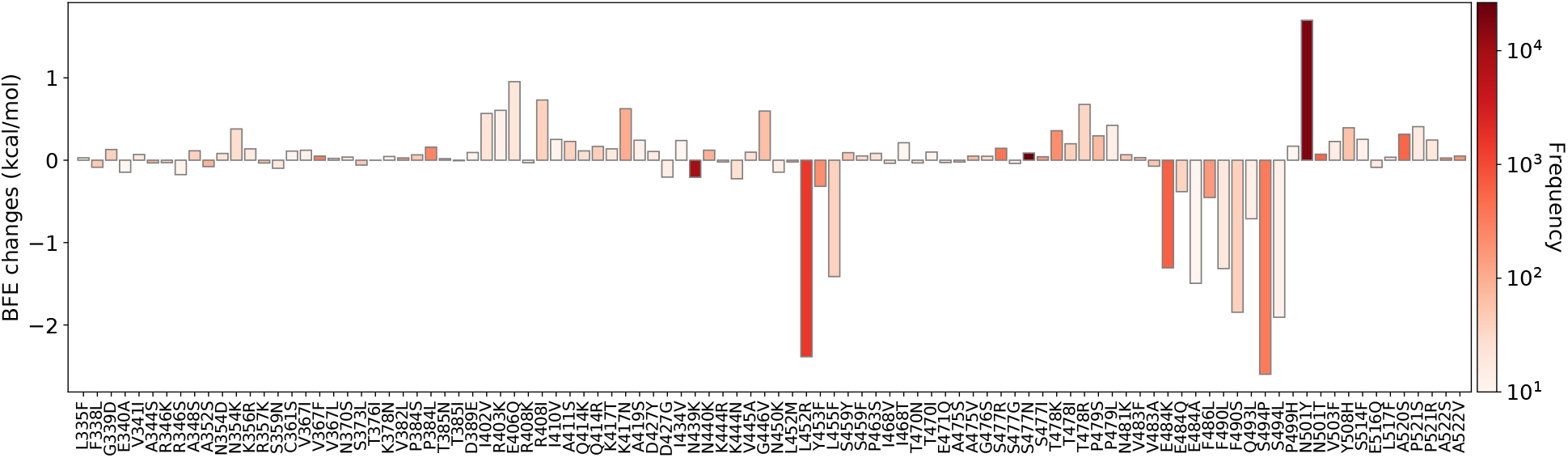
Illustration of SARS-CoV-2 mutation-induced binding free energy changes for the complexes of S protein and antibody CT-P59 (PDB: 7CM4). Here, mutations L452R, L455F, E484K/A, F490L/S, and S494P/L could potentially disrupt the binding of antibodies and S protein RBD.

#### 2.3.4 Antibodies C135 and C144

Lastly, we study C135 and C144, another antibody combination treatment currently on phase 1 study. Due to fact that there is no 3D structure of C135 and C144 binding to RBD simultaneously, we present their BFE change predictions based on PDB 7K8Z and 7K90, separately.

In the BFE change calculation of antibody C135, most mutations have mild BFE changes, while two mutations, R346K/S, induce large negative BFE changes, and three mutations, N440K, N450K, and P499H, lead to positive BFE changes greater than 0.5 kcal/mol. Notably, C135 is not an antibody that directly competes with ACE2 in terms of the binding domain. For mutations of emergent variants, K417T/N, L452N, and N501Y, they all have small BFE changes. With mild changes of most mutations, the antibody C135 could be a complement of other antibodies that are directly competing with ACE2 on the binding domain.

The last antibody is C144, which shares a part of the binding domain with ACE2. It is obvious that except for mutations E484K/Q/A, the rest mutations induce mild BFE changes. As the mutation E484K is part of the Brazil and South Africa variants, this antibody treatment could have antibody-escape effects. However, since most mutations lead to mild BFE changes and mutations K417N/T, L452R, T478K, and N501Y render mild positive BFE changes, this antibody can have neutralizing efficacy for many emerging variants, such as the UK, California, and Mexican variants.

## 3 Discussion

There are emerging variants spreading worldwide, which increase the virus transmissibility, reduce the neutralization of antibodies, and degrade the efficacy of antibody treatments or vaccines. The S protein plays the most important role in leading the virus to access the host cell. The RBD of S protein directly contacts ACE2, and its substitutions induced by variants can significantly weaken its binding with original antibodies that were either created from current vaccines or induced through existing antibody therapies. RBD mutations that enhance the RBD binding to ACE2 and weaken the RBD binding to many antibodies pose potential threats to vaccines and antibody therapies. Figure S1 in the Supporting information provides detailed analysis 514 RBD mutations to 16 antibodies.

**Figure 15:**
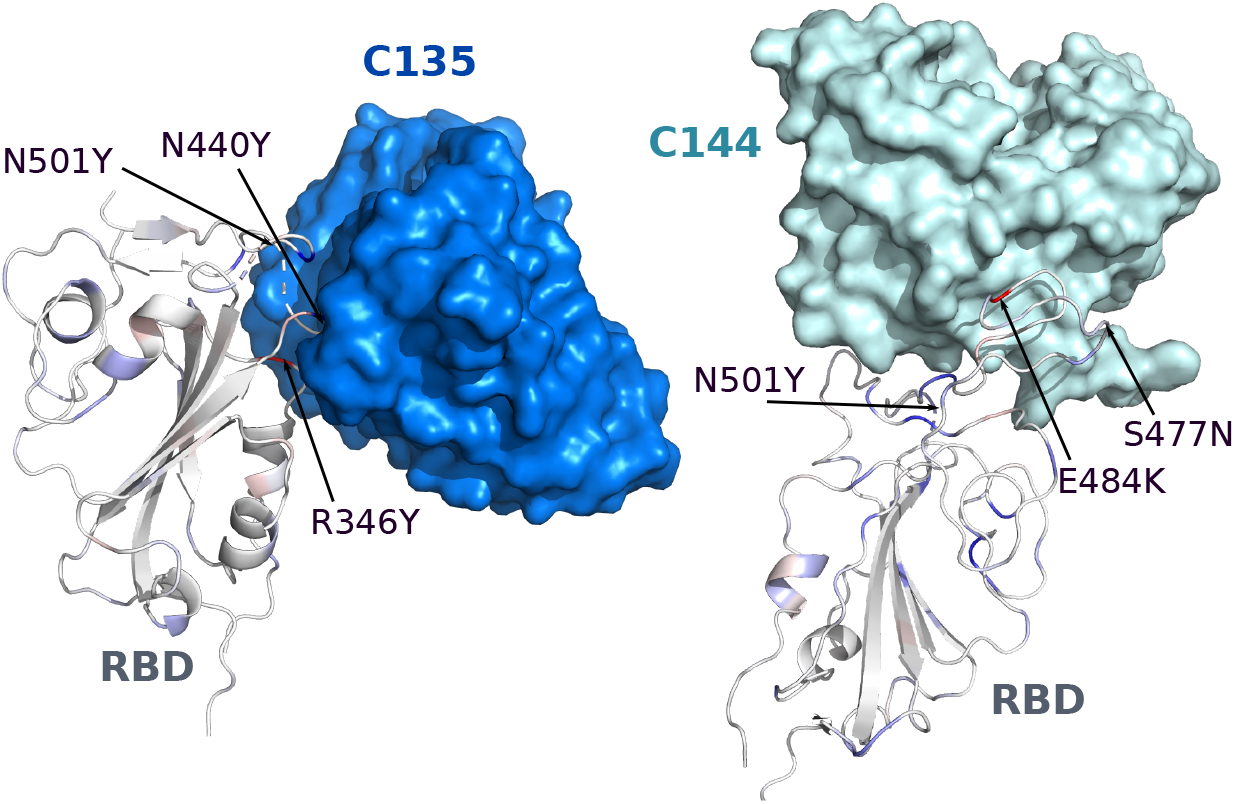
The binding complexes of S protein and antibodies C135 and C144.

**Figure 16:**
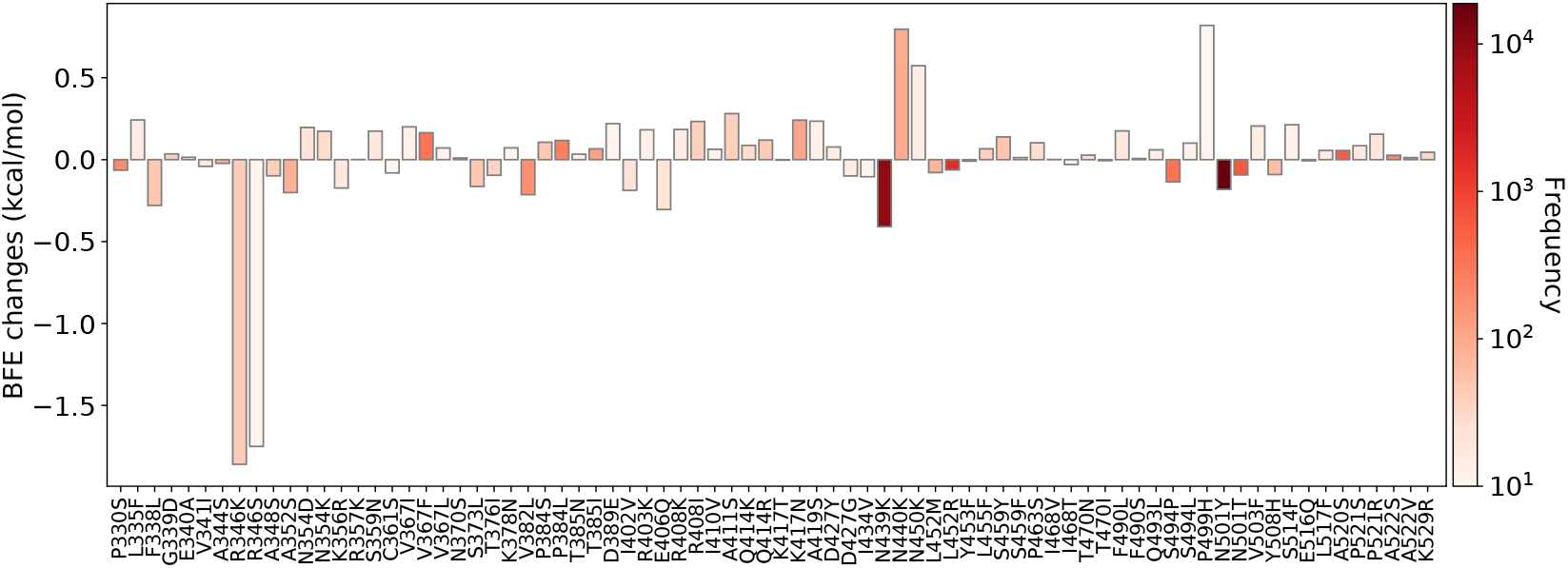
Illustration of SARS-CoV-2 mutation-induced binding free energy changes for the complexes of S protein and antibody C135 (PDB: 7K8Z). Here, mutations R346K/S could potentially disrupt the binding of antibodies and S protein RBD.

### B.1.1.7 lineage

The UK variant has one mutation, N501Y, on the RBD, which increases viral transmission [16] and severity based on hospitalizations [31]. However, for antibodies in clinical trials, it has a minor impact on neutralization in terms of BFE changes based on our predictions. Similar findings for the B.1.1.7 lineage have been reported for experimental neutralization by EUA therapeutics [17,18,32] and for other antibodies [10].

### P.1 lineage

The Brazil variant has three RBD mutations K417T, E484K, and N501Y. According to our BFE predictions, casirivimab (REGN10933) is moderately influenced by K417N and E484K on neutralization, while for imdevimab (RENG10987), the mutation impact is less significant. Regdanvimab (CT-P59) could still maintain its neutralizing capability. Bamlanivimab (LY-CoV555) shows a large fold reduction in susceptibility on mutation E484K in our prediction, which is consistent with a CDC report [18], while the combination of bamlanivimab and etesevimab gives a better response to P.1 lineage [17]. We hypothesize that CB6 is competing with LY-CoV555 and preserve its neutralization capacity with E484K. The combination of bamlanivimab and etesevimab has a large BFE reduction from P.1 lineage, which indicates that K417T has negative impacts on the binding.

**Figure 17:**
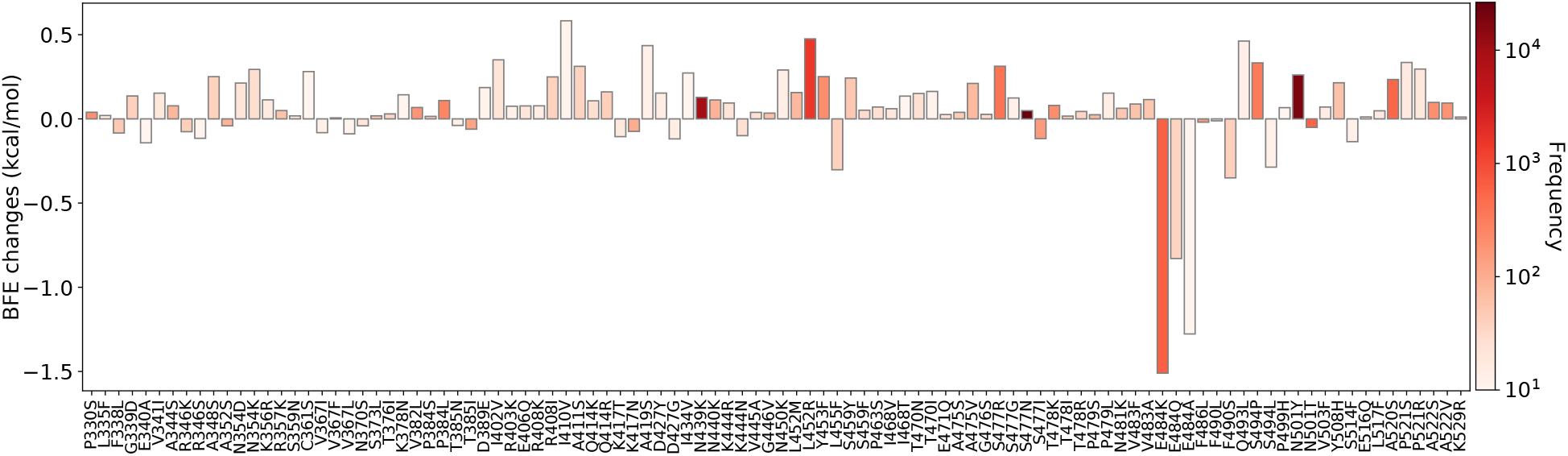
Illustration of SARS-CoV-2 mutation-induced binding free energy changes for the complexes of S protein and antibody C144 (PDB: 7K90). Here, mutations E484K/Q/A could potentially disrupt the binding of antibodies and S protein RBD.

### B.1.351 lineage

The South Africa variant is different from the Brazil variant only for one RBD mutation, i.e., K417N. We can claim a similar statement but moderate impacts on all the clinical trial antibodies. The same pattern can be found in the CDC report [17, 18] and the literature [10].

### B.1.427/429 lineage

For the California variants, the mutation, L452R, will have a negative impact on the neutralization for regdanvimab, but minimal impact on the neutralization by the two antibody combinations. L452R reduces the capacity of bamlanivimab, which can be shown by the prediction and the CDC report [18]. Interestingly, the fact of small impact on the combination, bamlanivimab and etesevimab, shown by the prediction and report [17] indicates that etesevimab dominants the binding process again.

### B.1.1.222 lineage

The Mexico variant involves RBD mutation T478K and has a larger positive BFE change on the binding of ACE2 and RBD. However, it has minor effects on existing antibodies.

### B.1.526 lineage

The New York variant is studied by only consider E484K. It reduces the neutralization of REGN10933, C144, and LY-CoV555. Based on our predictions, the impact on REGN10933 can be reduced if REGN10987 is in the treatment as well.

Figure 18 gives an overall comparison of experimental and predicted patterns of variant impacts on major antibody drug candidates. There is an excellent agreement between our predictions and various experimental data, except for a minor discrepancy. Specifically, our prediction shows a potential two-fold reduction in BFE for CB6 from B.1.1.7 due to N501Y (see Fig. 5), while the experiment records little change [10].

**Figure 18:**
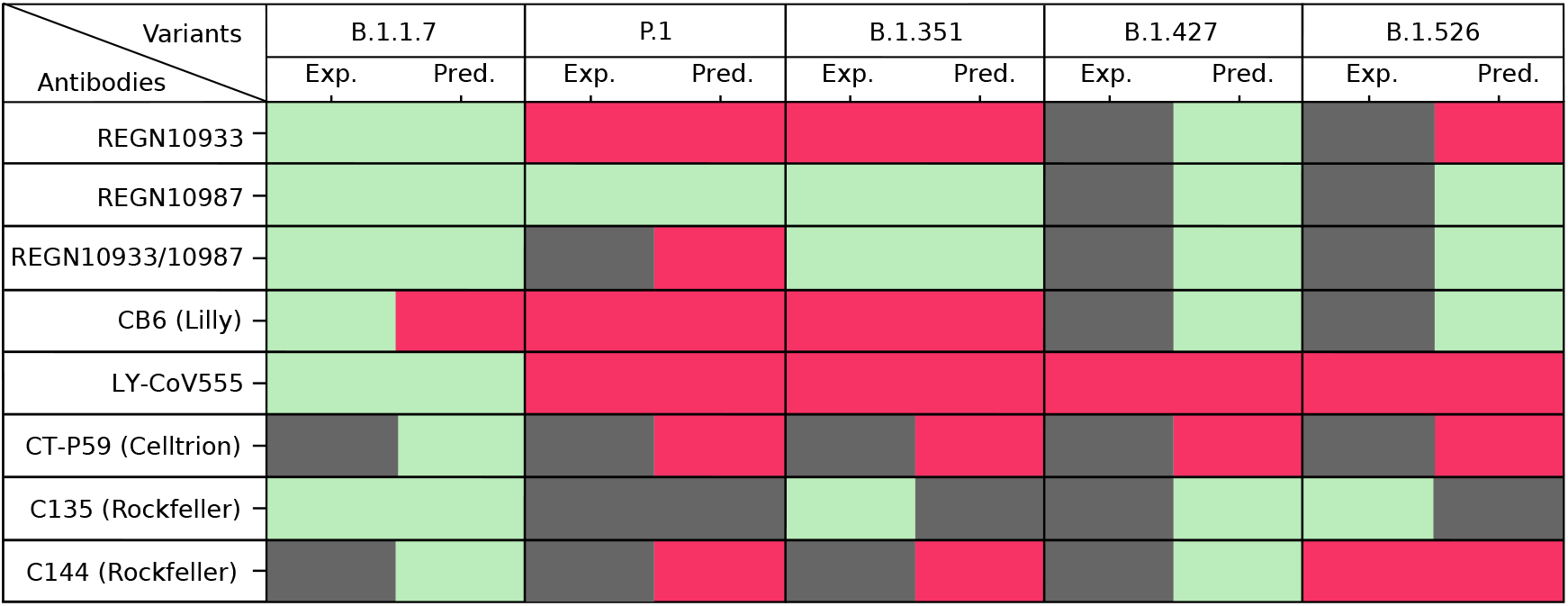
Comparison of experimental (Exp.) pattern and predicted (Pred.) pattern of the impact of SARS-CoV-2 variants on major antibody therapeutic candidates. Light green indicates mild or no change in neutralization; pink indicates significant reduction in neutralization; grey indicates no available data. RBD mutations in various variants: B.1.526: E484K; B.1.1.7: N501Y; B.1.427: L452R; P.1: K417T+E484K+N501Y; B.1.351: K417N+E484K+N501Y. The BFE changes are accumulated for multi-mutation predictions. Data resource: REGN10933 [10,30], REGN10987 [10,30], REGN10933/10987 [10], CB6 [10,30], C135 [10,33], C144 [33], and LY-CoV555 [10,18]

In summary, the Eli Lilly antibody therapies bamlanivimab and etesevimab are likely compromised by known emerging variants and other high-frequency mutations V483F/A, E484Q/V/A/G/D, F486L, F490L/V/S, Q493L, and S494P. For Regeneron antibody therapies casirivimab and imdevimab, while there is no experimental data regarding K417T from variant P.1, our predictions indicate that there is a potential compromise from mutation K417T. Additionally, Regeneron antibodies are prone to high-frequency mutations N439K, G446V, E484G, and F486L. The Celltrion antibody therapy regdanvimab (i.e., CT-P59) is predicted to be compromised by variants P.1, B1.351, B.1.427, and B.1.526, although there is no experimental data now. It can also be weakened by high-frequency mutations L455F, E484A, F490L/S, and S494P/L. Rockefeller University antibody C135 can be evaded by high-frequency mutations R346K/S. The antibody C144 from Rockefeller University is prone to variants P.1, B1.351, and B.1.526, while the experiment has only confirmed the adversarial impact of variant B.1.526. Additionally, it can be compromised by high-frequency mutations E484Q/A. In the Supporting information, we further identify that low-frequency RBD mutations V401I/L, I402V, E406G, Q409L, I410V, D420A/G, N422S, N448D, N450D, Y453F, F456L, Y473F, E484Q/A/G/D, G485S/R/C/V, F486L/V/C, F490I/L/V/Y/S, S393A/L, N501I, and Y508S have potential to become future vaccine or antibody escape variants. These mutations are predicted to enhance the RBD binding to ACE2 while weaken the binding between RBD and most antibodies.

## 4 Methods

### 4.1 Genome sequence data and pre-processing

Complete SARS-CoV-2 genome sequences are available from the GISAID database [34]. In this work, a total of 261,348 complete SARS-CoV-2 genome sequences with high coverage and exact collection date are downloaded from the GISAID database [34] (https://www.gisaid.org/) as of March 10, 2021. We take the first complete SARS-CoV-2 genome from the GenBank (NC_045512.2) as the reference genome [35], and the multiple sequence alignment is applied by the Clustal Omega [36, 37] with default parameters, which results in 27,530 single nucleotide polymorphism profiles. On the S protein RBD, i.e., residues 329 to 530, 514 non-degenerate mutations are found. Among them, 95 mutations have been observed for more than 10 times.

### 4.2 Machine learning datasets

Dataset is important to train accurate machine learning models. Both the BFE changes and enrichment ratios describe the effects on the binding affinity of protein-protein interactions. Therefore, integrating both kinds of datasets can improve the prediction accuracy. Especially, due to the urgency of COVID-19, the BFE changes of SARS-CoV-2 data are rarely reported, while the enrichment ratio data via high-throughput deep mutations are relatively easy to obtain. The most important dataset that provides the information for binding free energy changes upon mutations is the SKEMPI 2.0 dataset [38]. The SKEMPI 2.0 is an updated version of the SKEMPI database, which contains new mutations and data from other three databases: AB-Bind [39], PROXiMATE [40], and dbMPIKT [41]. There are 7,085 elements, including single- and multi-point mutations in SKEMPI 2.0. 4,169 variants in 319 different protein complexes are filtered as single-point mutations are used for our TopNetTree model training. Moreover, SARS-CoV-2 related datasets are also included to improve the prediction accuracy after a label transformation. They are all deep mutation enrichment ratio data, mutational scanning data of ACE2 binding to the receptor-binding domain (RBD) of the S protein [42], mutational scanning data of RBD binding to ACE2 [43, 44], and mutational scanning data of RBD binding to CTC-445.2 and of CTC-445.2 binding to the RBD [44]. Note that our training datasets used in the validation do not include the test dataset, which is a mutational scanning data of RBD binding to ACE2.

### 4.3 Feature generation of PPIs

Algebraic topology [45, 46] has had tremendous success in describing biochemical and biophysical properties [47]. Element-specific and site-specific persistent homology can effectively simplify the structural complexity of protein-protein complex and extract the abstract properties of the vital biological information in PPIs [6, 48]. The algebraic topological analysis on PPIs is constructed based on a series of atom subsets of complex structures, which are atoms of the mutation sites, 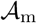, atoms in the neighborhood of the mutation site within a cut-off distance *r*, 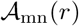, antibody atoms within *r* of the binding site, 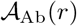, antigen atoms within *r* of the binding site, 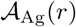, and atoms in the system that has atoms of element type of {C, N, O}, 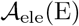. Additionally, a bipartition graph is introduced to describe the antibody and antigen in PPIs. Then, molecular atoms construct point clouds for simplicial complex, which is a finite collection of sets of linear combinations of points. We apply the Vietoris-Rips (VR) complex for dimension 0 topology, and alpha complex for point cloud of dimensions 1 and 2 topology [47]. Overall, element-specific and site-specific persistent homology is devised to capture the multiscale topological information over different scales along a filtration [45] and is important for our machine learning predictions.

#### 4.3.1 Simplex and simplicial complex

Given a set of independent *k* +1 points *U* = {*u*_0_, *u*_1_,…, *u_k_*} in 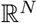, the convex combination is a point 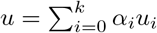, where 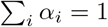 and *α_i_* ≥ 0. The convex hull of *U* is the collection of convex combinations of *U*, and a *k*-simplex *σ* is the convex hull of *k*+1 independent points *U*. For example, a 0-simplex is a point, a 1-simplex is an edge, a 2-simplex is a triangle, and a 3-simplex is a tetrahedron. A proper *m*-face of the *k*-simplex is a subset of the *k* + 1 vertices of a *k*-simplex with *m* +1 vertices forms a convex hull in a lower dimension and *m* < *k*. The boundary of a *k*-simplex *σ* is defined as a sum of all its (*k* – 1)-faces as

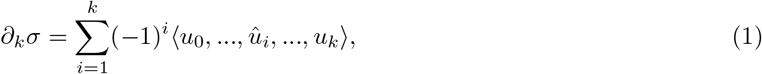

where 〈*u*_0_,…, *û_i_,…, u_k_*〉 is a convex hull formed by vertices of *σ* excluding *u_i_*. A simplicial complex denotes by *K* is a collection of finitely many simplices forms a simplicial complex. Thus, faces of any simplex in *K* are also simplices in *K*, and intersections of any 2 simplices are only faces of both or an empty set. A *k*-simplex *σ* = (*u*_*i*_0__,…, *u_i_k__*) is in Vietoris-Rips complex *R^r^*(*U*) if and only if 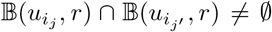 for *j, j′* ∈ [0, *k*] and is in alpha complex *A^r^*(*U*) if and only if 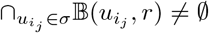.

#### 4.3.2 Homology

For a simplicial complex *K*, a *k*-chain *c_k_* of *K* is a formal sum of the *k*-simplices in *K* defined as 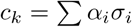, where *σ_i_* is the *k*-simplices and *α_i_* is coefficients. *α_i_* can be in different fields such as 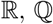, and 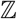. Typically, *α_i_* is chosen to be 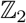, which is {−1, 0, 1} and forms an Abelian group 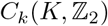. Then, the boundary operator can be extended to a *k*-chain *c_k_* as

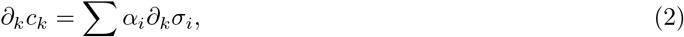

such that *∂_k_*: *C_k_* → *C*_*k*−1_ and satisfies 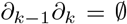, follows from that boundaries are boundaryless. The chain complex is defined as a sequence of complexes by boundary maps is called a chain complex

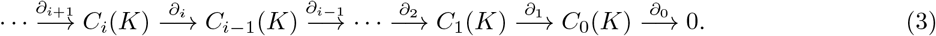

The *k*-homology group is the quotient group defined by taking *k*-cycle group module of *k*-boundary group as

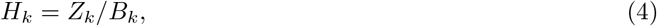

where *H_k_* is the *k*-homology group, and *k*-cycle group *Z_k_* and the *k*-boundary group *B_k_* are the subgroups of *C_k_* defined as,

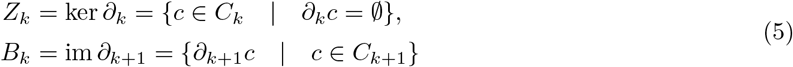

The Betti numbers are defined by the ranks of *k*th homology group *H_k_* as *β_k_* = rank(*H_k_*). *β*_0_ reflects the number of connected components, *β*_1_ reflects the number of loops, and *β*_2_ reflects the number of cavities.

#### 4.3.3 Filtration and persistent homology

A filtration of a topology space *K* is a nested sequence of *K* such that

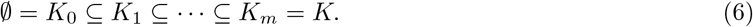

Then, a sequence of chain complexes and a homology sequence are constructed on the filtration. The *p*th persistent of *k*th homology group of *K_t_* are defined as

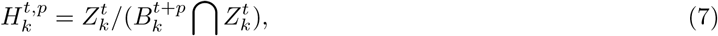

and the Betti numbers 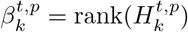. These persistent Betti numbers are applied to represent topological fingerprints.

#### 4.3.4 Auxiliary features

Features of topological invariants are not enough to reflect the whole picture of PPIs. Importantly, chemical and physical information, including surface areas, partial charges, Coulomb interactions, van der Waals interaction, electrostatic solvation free energy, mutation site neighborhood amino acid composition, pKa shifts, and secondary structure, is added as auxiliary features to improve the predictive power of the machine learning model [5].

### 4.4 Machine learning and deep learning algorithms

We illustrate the construction of a topology-based network (TopNet) model for the BFE change prediction of protein-protein interactions (PPIs) on SARS-CoV-2 studying. These approaches have been widely applied in studying protein-ligand and protein-protein binding free energy predictions [5, 6]. Firstly, one ensemble method, gradient boosting decision tree (GBDT), is studied as baselines in comparison to deep neural network methods. The ensemble methods naturally handle correlation between descriptors and are robust to redundant features. Therefore, they usually do not depend on a sophisticated feature selection procedure and a complicated grid search of hyper-parameters. The implemented GBDT is a function from the scikit-learn package (version 0.22.2.post1) [49]. The number of estimators and the learning is optimized for ensemble methods as 20000 and 0.01, respectively. For each set, 10 runs (with different random seeds) were done and the average result is reported in this work. Considering a large number of features, the maximum number of features to consider is set to the square root of the given descriptor length for GBDT methods to accelerate the training process. The parameter setting shows that the performance of the average of sufficient runs is decent.

A neural network is a network of neurons that maps an input feature layer to an output layer. The neural network simulates a biological brain solves problems with numerous neuron units by backpropagation to update weights on each layer. To reveal the facts of input features at different levels and abstract more properties, one can construct more layers and more neurons in each layer, which is known as a deep neural network. Optimization methods for feedforward neural networks and dropout methods are applied to prevent overfitting.

#### 4.4.1 Optimization

To train feedforward neural networks, backpropagation is applied where the loss function is evaluated at the output layer and is propagated backward through the network to update the model’s weights and bias. As the calculation of gradient required, one popular approach is the stochastic gradient descent (SGD) method with momentum which estimates a small portion of training data and applies the idea of exponentially weighted averages. Thus, the momentum term can accelerate the convergence of the algorithm. A popular way to implement the SGD with momentum is given as

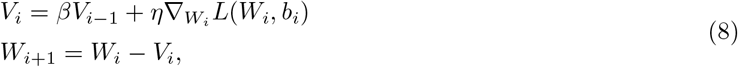

where *W_i_* is the parameters in the network, *L*(*W_i_, b_i_*) is the objective function, *η* is the learning rate, *X* and *y* are the input and target of the training set, and *β* ∈ [0,1] is a scalar coefficient for the momentum term.

#### 4.4.2 Dropout

Fully connected layers possess a large number of degrees of freedom. This can easily cause an overfitting issue, while the dropout technique is an easy way of preventing network overfitting. [ref] In the training process, hidden units are randomly set zero values to their connected neurons in the next layer. Suppose that a percentage of neurons at a certain layer is chosen to be dropped during training. The number of computed neurons of this layer is equal to the neuron number multiplied by a coefficient such as 1-*p*, where *p* is the dropout rate. Then, in the testing process, the output of these layers is computed by randomly dropouts the same rate of neurons, to approximate the network in each training step.

#### 4.4.3 Deep learning algorithms

A deep neural network (DNN) consists of multi-layers of neurons. In the output layer, the single neuron gets full connections with the last hidden layer and calculates predictions. Notice that the network is constructed for mutation-induced BFE changes, one should preserve the consistency of all labels. An optimizer is used to minimize the following loss function:

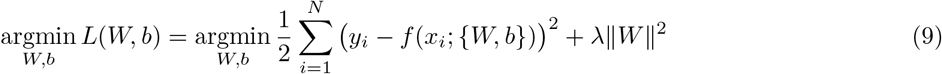

where *N* is the number of samples, *f* is a function of the feature vector *x_i_* parametrized by a weight vector *W* and bias term *b*, and λ represents a penalty constant.

### 4.5 Validation

Here, we present a validation of our BFE change prediction for mutations on S protein RBD compared to the experimental deep mutations enrichment data [44]. Figure 19 presents a comparison between experimental deep mutations enrichment data and BFE change predictions on SARS-CoV-2 RBD binding to ACE2. In the heatmap of Figure 19, it is obvious that the predicted BFE changes are highly correlated to the enrichment ratio data. Both BFE changes and enrichment ratios describe the affinity changes of the S protein RBD-ACE2 complex induced by mutations.

**Figure 19:**
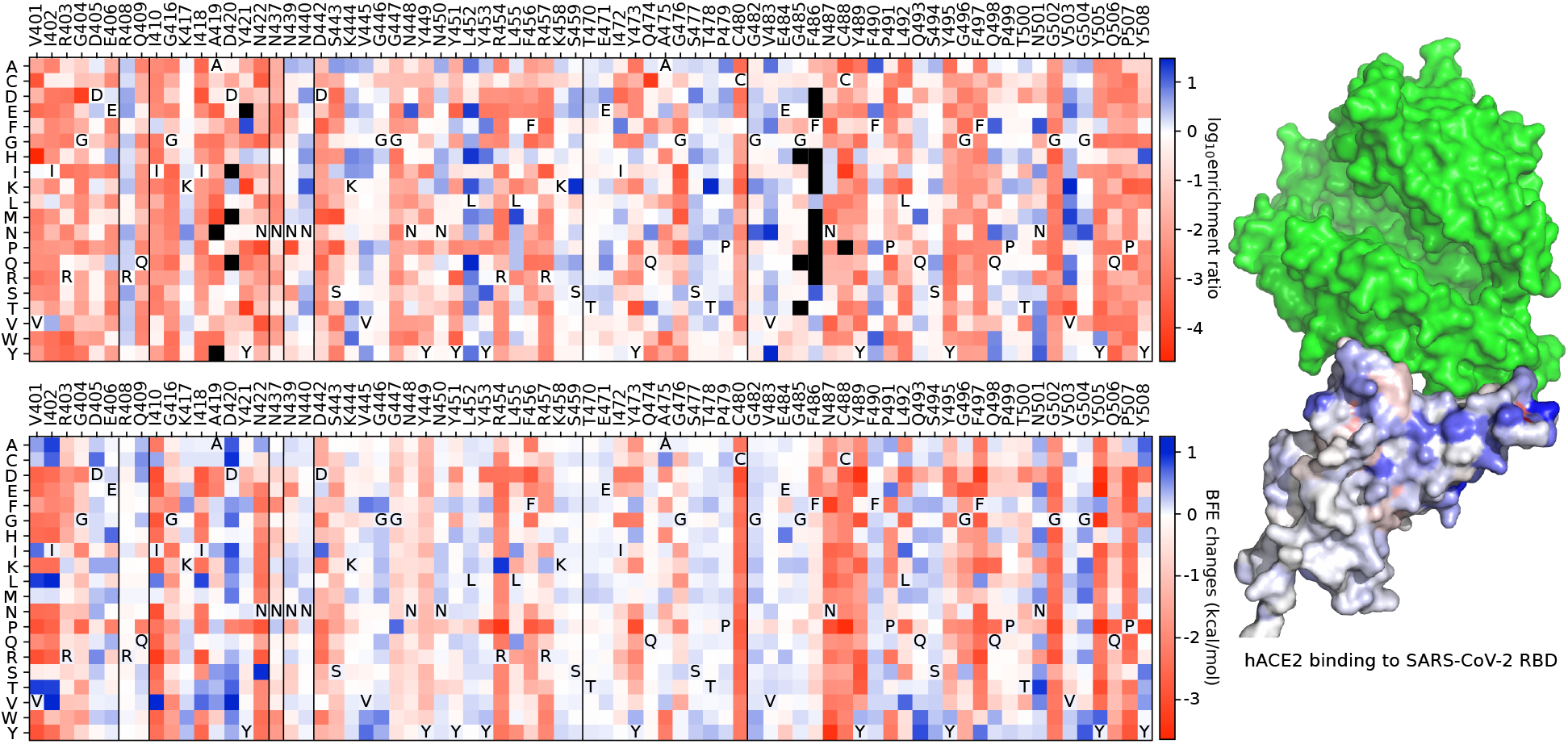
A comparison between experimental RBD deep mutation enrichment data and predicted BFE changes for SARS-CoV-2 RBD binding to ACE2 (6M0J) [44]. **Top left**: deep mutational scanning heatmap showing the average effect on the enrichment for single-site mutants of RBD when assayed by yeast display for binding to the S protein RBD [44]. **Right**: RBD colored by average enrichment at each residue position bound to the S protein RBD. **Bottom left**: machine learning predicted BFE changes for single-site mutants of the S protein RBD.

## Data and model availability

The SARS-CoV-2 single nucleotide polymorphism data in the world is available at [Mutation Tracker](https://users.math.msu.edu/users/weig/SARS-CoV-2_Mutation_Tracker.html). The machine learning training datasets and the trained machine learning model are available at [TopNetmAb](https://github.com/WeilabMSU/TopNetmAb). The related training process is described in Supporting information.

## Supporting information

The supporting information is available for S1 BFE changes for the complexes of S protein RBD binding to antibodies or ACE2 induced by 514 RBD mutations 2 and S2 Machine learning models.

## Acknowledgment

This work was supported in part by NIH grant GM126189, NSF grants DMS-2052983, DMS-1761320, and ΠS-1900473, NASA grant 80NSSC21M0023, Michigan Economic Development Corporation, Bristol-Myers Squibb 65109, and Pfizer. The authors thank The IBM TJ Watson Research Center, The COVID-19 High Performance Computing Consortium, NVIDIA, and MSU HPCC for computational assistance. RW thanks Dr. Changchuan Yin for useful discussion.

