## Supplementary Material for "Revealing the threat of emerging SARS-CoV-2 mutations to antibody therapies"

Jiahui Chen[1], Kaifu Gao[1], Rui Wang[1], and Guo-Wei Wei[1,2,3*]
[1] Department of Mathematics,
Michigan State University, MI 48824, USA.
[2] Department of Electrical and Computer Engineering,
Michigan State University, MI 48824, USA.
[3] Department of Biochemistry and Molecular Biology,
Michigan State University, MI 48824, USA.
[†]First three authors contributed equally.

April 12, 2021

#### S1 BFE changes for the complexes of S protein RBD binding to antibodies or ACE2 induced by 514 RBD mutations

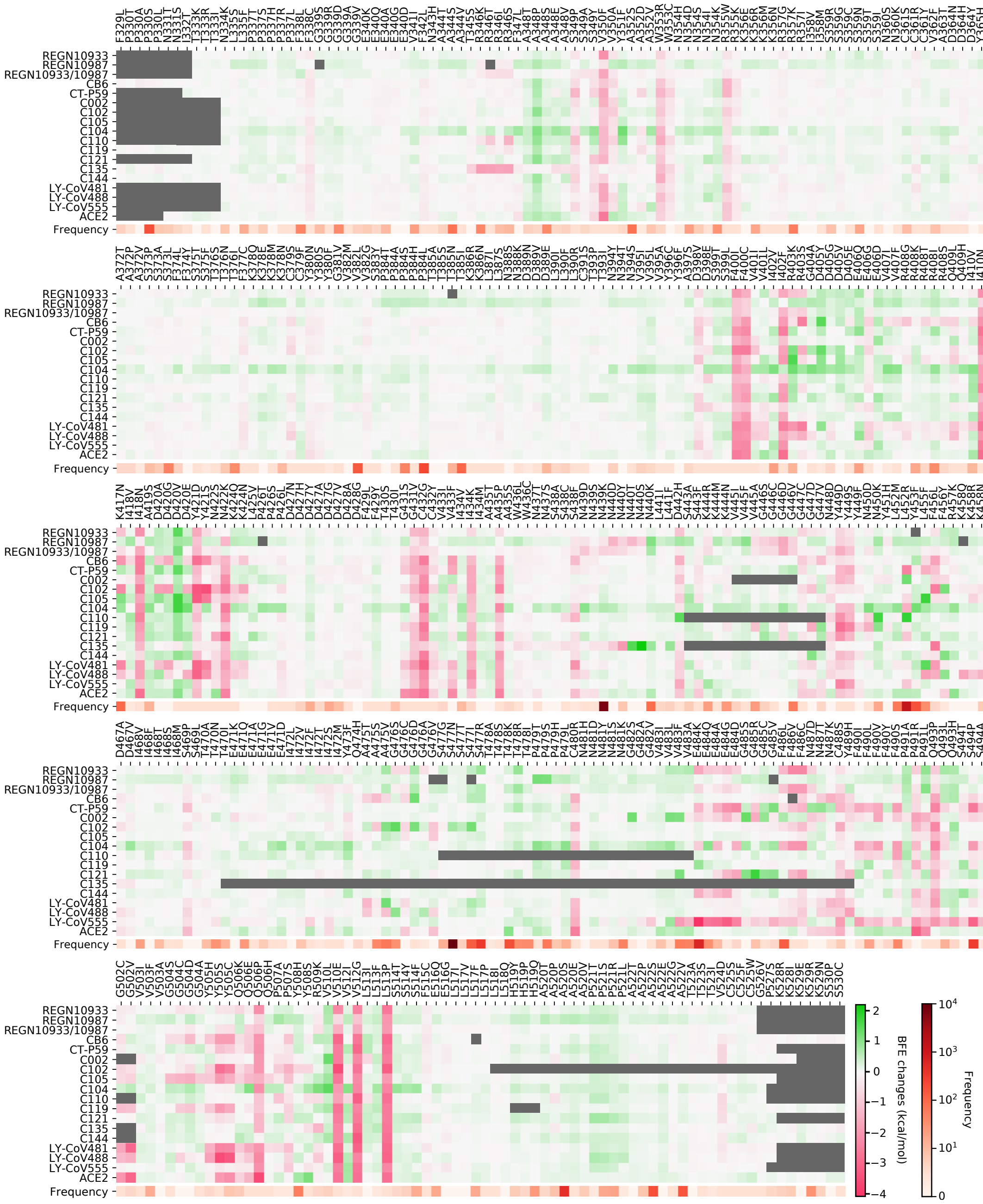

Figure S1: The heatmap of BFE changes for the complexes of S protein RBD binding to antibodies or ACE2 induced by 514 RBD mutations. Mutation frequencies are also given. The gray color indicates no data available due to incomplete structures.

In Figure S1, we present the heatmap of BFE changes for the complexes of S protein RBD binding to antibodies or ACE2 induced by 514 known RBD mutations. We also provide mutation frequencies, i.e., the numbers of observed times, for various mutations. It can be seen that the BFE changes of the RBD-ACE2 complex are highly correlated with the frequencies of mutations. High-frequency mutations tend to have positive BFE changes due to natural selection. These are the so-called fast-growing mutations. These mutations can be disruptive to vaccines and antibody therapies if they induce significant negative BFE changes to the RBD binding of many antibodies. Therefore, in Figure S1, mutations V401I/L, I402V, E406G,

Q409L, I410V, D420A/G, N422S, N448D, N450D, Y453F, F456L, Y473F, E484Q/A/G/D, G485S/R/C/V, F486L/V/C, F490I/L/V/Y/S, S393A/L, N501I, and Y508S can become antibody-escape mutations. Another class of fast-growing mutations leads to mostly positive BFE changes to antibodies, but more positive BFE changes to ACE2, such as V483I/L/F. This type of mutation might not evade vaccines and antibody therapies but be more transmissible like the Mexico variant T478K. Some of the RBD mutations are predicted to significantly weaken the RBD binding to both ACE2 and all antibodies. However, it will be unlikely for them to disrupt antibody therapies and vaccines due to natural selection. There are still RBD mutations that weaken the RBD binding to ACE2 but produce positive BFE changes to antibodies, such as Q498K. As discussed above, this type of mutation would not happen very often in a population.

### S2 Machine learning models

Two machine learning models, gradient boosting decision tree (GBDT) and artificial neural network (ANN) are implemented, where GBDT is a baseline model for the comparison to ANN.

#### S2.1 Auxiliary features

We discuss the detail of element-specific and site-specific persistent homology in the main content but have a glimpse of the auxiliary feature generation. However, these auxiliary features, namely other chemical and physical information that has not been incorporated into element-specific persistent homology, can improve the predictive power of our machine learning models. These features are concatenated with topological features for GBDT training and ANN training. Auxiliary features are categorized into residue-level and atom-level ones.

##### S2.1.1 Residue-level features

**S2.1.1.1 Mutation site neighborhood amino acid composition** Neighbor residues are the residues within 10 Å of the mutation site. Distances between residues are calculated based on residue $C_\alpha$ atoms. Six categories of amino acid residues are counted, which are hydrophobic, polar, positively charged, negatively charged, special cases, and pharmacophore changes. The count and percentage of the 6 amino acid groups in the neighbor site are regrading as the environment composition features of the mutation site. The sum, average, and variance of residue volumes, surface areas, weights, and hydropathy scores are used but only the sum of charges is included.

**S2.1.1.2 pKa shifts** The pKa values are calculated by the PROPKA software [2], namely the values of 7 ionizable amino acids, namely, ASP, GLU, ARG, LYS, HIS, CYS, and TYR. The maximum, minimum, sum, the sum of absolute values, and the minimum of the absolute value of total pKa shifts are calculated. We also consider the difference of pKa values between a wild type and its mutant. Additionally, the sum and the sum of the absolute value of pKa shifts based on ionizable amino acid groups are included.

**S2.1.1.3 Position-specific scoring matrix (PSSM)** Features are computed from the conservation scores in the position-specific scoring matrix of the mutation site for the wild type and the mutant as well as their difference. The conservation scores are generated by PSI-BLAST [1].

**S2.1.1.4 Secondary structure** The SPIDER2 software is used to compute the probability scores for residue torsion angle and residues being in a coil, alpha helix, and beta strand based on the sequences for the wild type and the mutant [13].

##### S2.1.2 Atom-level features

Seven groups of atom types, including C, N, O, S, H, all heavy atoms, and all atoms, are considered when generating the element-type features. Meanwhile, other three atom types, i.e., mutation site atoms, all heavy atoms, and all atoms, are used when generating the general atom-level features.

**S2.1.2.1 Surface areas** Atom-level solvent excluded surface areas are computed by ESES [8].

**S2.1.2.2 Partial changes** Partial change of each atom is generated by pdb2pqr software [6] using the Amber force field [4] for wild type and CHARMM force field [3] for mutant. The sum of the partial charges and the sum of absolute values of partial charges for each atomic group are collected.

**S2.1.2.3 Atomic pairwise interaction interactions** Coulomb energy of the $i$th single atom is calculated as the sum of pairwise coulomb energy with every other atom as

$$C_i = \sum_{j,j\neq i} k_e \frac{q_i q_j}{r_{ij}}, \tag{1}$$

where $k_e$ is the Coulomb's constant, $r_{ij}$ is the distance of $i$th atom to $j$th atom, and $q_i$ is the charge of $i$th atom. The van der Waals energy of the $i$th atom is modeled as the sum of pairwise Lennard-Jones potentials with other atoms as

$$V_i = \sum_{j,j\neq i} \epsilon \Big[ \big(\frac{r_i + r_j}{r_{ij}}\big)^{12} - 2\big(\frac{r_i + r_j}{r_{ij}}\big)^{6} \Big], \tag{2}$$

where $\epsilon$ is the depth of the potential well, and $r_i$ is van der Waals radii.

In atomic pairwise interaction, 5 groups (C, N, O, S, and all heavy atoms) are counted both for Coulomb interaction energy and van der Waals interaction energy.

**S2.1.2.4 Electrostatic solvation free energy** Electrostatic solvation free energy of each atom is calculated using the Poisson-Boltzmann equation via MIBPB [5] and are summed up by atom groups.

#### S2.2 Training process description

For the mentioned databases in main content, crystal structures of the wild type binding free energy changes are given at the first place. Then crystal structures of mutant are generated by scap utility in the Jackal package [12] which predicts side-chain conformations on a given backbone. Missing residues are not made up, only missing atoms are fixed. VMD is applied to select the structures of mutation site and binding site [7].

##### S2.2.1 Topology features

Element-specific persistent homology $H_0$ features are generated by the Vietoris-Rips complex barcodes for the distances of filtration from 2 Å to 11 Å with the interval length of 1 Å. Element-specific persistent homology $H_1$ and $H_2$ barcodes are constructed by alpha shape with cutoff value equal to 12 Å. For each barcode of $H_1$ and $H_2$, the statistical values, namely sum, min, max, mean, and standard deviation are collected. The topological features are generated by GUDHI [11].

##### S2.2.2 TopGBDT: Topology based GBDT model

The hyperparameters for TopGBDT are generated by gird searches based on 10-fold cross validation. The hyperparameters are set to be `n_estimators = 20000`, `max_depth = 7`, `learning_rate = 0.01`, `criterion = friedman_mse`, `subsample = 0.6`, `max_features = sqrt`, `min_samples_split=3`, and `loss=ls`. The implemented GBDT is a function from the scikit-learn package (version 0.22.2.post1) [10].

###### S2.2.3 TopANN: Topology based ANN model

The network layers and the number of neurons in each layer are found by gird searches based on a 10-fold cross validation. Then the hyperparameters of stochastic gradient descent (SGD) with momentum are set up based on the network structure. The network has 7 layers with 8000 neurons in each layer. For SGD with momentum, the hyperparameters are `momentum = 0.9` and `weight_decay=0`. The implemented ANN is based on Pytorch [9].
